# Reconstructing donor genotypes from scRNA-seq for downstream eQTL and HLA–TCR analysis

**DOI:** 10.64898/2026.09.30.755703

**Authors:** Hannah Kockelbergh, Jennifer Astley, Katherine M. Kichula, Zhipeng Zhang, Esther S. Ng, COvid-19 Multi-omics Blood ATlas (COMBAT) Consortium, Calliope A. Dendrou, Amanda Y. Chong, Robert A. Watson, Benjamin P. Fairfax, Julian C. Knight, Gavin Band, Paul J. Norman, Mary N. Carrington, Stephen N. Sansom, Alexander J. Mentzer, Luke Jostins-Dean, Yang Luo

## Abstract

Single-cell eQTL studies can reveal heterogeneous genetic effects on gene expression that vary with cell type, but their wider application has been constrained by the frequent absence of matched genotype data. Here, we present single-cell Transcriptome Allelic Prediction and Association Studies (scTAPAS), a bioinformatic framework for reconstructing donor genotypes directly from single-cell RNA-sequencing reads and downstream genetic analysis. Applying scTAPAS to 5’ single-cell RNA-sequencing data from the COVID-19 Multi-omic Blood Atlas (COMBAT), we show that reference-based imputation of sparse RNA-derived genetic information generates genotype dosages sufficiently accurate for association testing. Despite testing only 18.0% of the variants available from conventional array-based genotyping and imputation, scTAPAS recovers 68.6% of cell type-eGene pairs identified using conventional array-based genotypes, with strong concordance in effect sizes. We further show that scTAPAS-derived variants can support imputation of classical HLA alleles and enable HLA-aware QTL analyses, including associations between HLA variation and TCR repertoire at single-cell resolution. Using matched single-cell TCR-sequencing data, we identify CD4+ T cell-specific associations between HLA class II alleles and T-cell receptor (TCR) Vα gene segment usage, highlighting the potential of analyses that complement TCR sequencing at single-cell resolution with HLA types. These results show that genetic information recovered from single-cell RNA sequencing data can support both regulatory and immunogenetic association analyses, expanding the discovery potential of single-cell cohorts for which conventional genotyping or HLA typing is unavailable.

## Introduction

Single-cell RNA sequencing (scRNA-seq) enables the identification of expression quantitative trait loci (eQTLs) within individual cell types, providing insights into genetic regulation that are obscured in bulk transcriptomic analyses [1], [2], [3], [4], [5], [6]. By resolving cellular heterogeneity, single-cell eQTL (sc-eQTL) studies have the potential to uncover cell type-specific regulatory effects and improve our understanding of how genetic variation influences cellular function in health and disease. However, broader application of sc-eQTL analysis has been constrained by the limited availability of scRNA-seq cohorts with matched genotype data. Although genetic variants can be detected directly from scRNA-seq reads, RNA-derived variant information is sparse and unevenly distributed across the genome because of allelic dropout and expression bias. As a result, while scRNA-seq-derived variant calling alone is sufficient for applications such as donor demultiplexing [7], [8], [9], [10], [11], it does not generally provide sufficiently dense and accurate genotypes to support robust genetic analyses.

We propose genotype imputation to recover donor genotypes directly from scRNA-seq reads, overcoming the limitations of direct variant calling. By integrating sparse variant information from RNA reads with population reference panels, we reconstruct accurate genotype dosages suitable for association testing. This strategy has been applied to low-coverage DNA sequencing [12], [13], but its utility for reconstructing donor genotypes from scRNA-seq and supporting single-cell genetic analyses remains unexplored. Recovering donor genetic information from scRNA-seq could also enable analyses beyond conventional eQTL mapping. This is particularly relevant for single-cell studies that combine transcriptome profiling with T-cell receptor (TCR) sequence. TCR repertoires are shaped by human leukocyte antigen (HLA) variation, as HLA molecules determine the peptide repertoire presented to T cells during thymic selection and immune responses. However, HLA types are also frequently unavailable in single-cell cohorts. Inferring HLA alleles from the same scRNA-seq data used for cellular profiling would therefore expand the range of immunogenetic analyses possible in existing single-cell studies.

Here, we develop single-cell Transcriptome-derived Allele Prediction and Association Studies (scTAPAS), a framework for reconstructing donor genotypes and enabling eQTL analysis from scRNA-seq alone. Using a deeply phenotyped single-cell cohort with matched array-based genotypes, HLA sequencing and single-cell TCR sequencing [14], we benchmark scTAPAS-derived genotypes against conventional genotyping and imputation and evaluate their performance in cell type-resolved cis-eQTL mapping. We then extend this strategy to the HLA genes, assessing whether RNA-derived genetic information can support classical HLA imputation and HLA-TCR association analysis.

## Results

### Method overview

We developed scTAPAS to infer donor-level genetic variation from scRNA-seq by combining donor-level read aggregation with reference-based imputation. We first consolidated all sequencing reads aligned to chromosome 6 into donor-specific alignment files that retain genetic information distributed across the transcriptome (**Fig. 1a**). We then processed these per-individual sequencing files with QUILT2 [13], [15], a read-aware imputation method that uses sparse low-coverage sequencing reads and large haplotype reference panels to infer genome-wide genotype dosages (**Fig. 1b**). Following variant-level quality control, scTAPAS-derived genotypes can be used directly for cell type-resolved *cis*-eQTL mapping (**Fig. 1c**).

**Fig 1.**
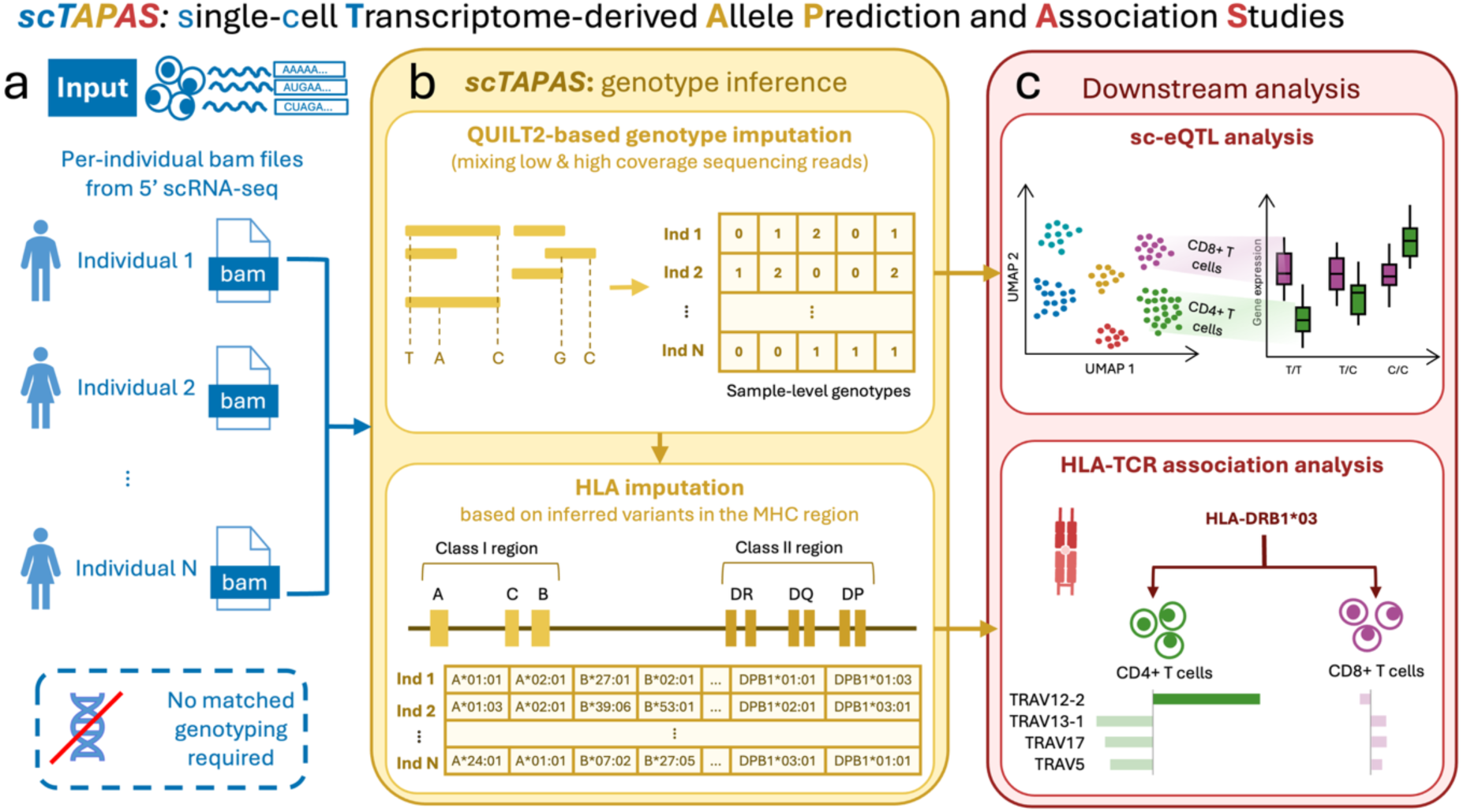
Overview of the scTAPAS workflow. scTAPAS aggregates scRNA-seq reads across cells within each donor (**a**) and applies QUILT2 to infer genotypes from sparse RNA-derived genetic information. Variants in the MHC region may then be used to impute classical HLA alleles and amino acid polymorphisms (**b**). Quality-controlled variants are used for downstream genetic analyses, including *cis*-eQTL mapping and single-cell HLA-TCR association studies (**c**).

To extend the workflow to immunogenetic analyses, scTAPAS uses inferred variants within the major histocompatibility complex (MHC) region to impute classical HLA alleles and amino acid polymorphisms (**Fig. 1b**). HLA features can then be integrated with matched single-cell TCR sequencing data to test associations between HLA variation and TCR repertoire features across T-cell populations (**Fig. 1c**). Thus, scTAPAS provides a common starting point for both cell type-resolved regulatory analyses and HLA-focused immunogenetic analyses using genetic information recovered from scRNA-seq.

### scTAPAS infers donor genotypes from scRNA-seq with high concordance

To evaluate scTAPAS, we applied the method to a previously published 5’ scRNA-seq cohort from the COVID-19 Multi-omic Blood Atlas (COMBAT) [5], [14]. Briefly, COMBAT profiles cellular states in blood over the course of infectious diseases at varying severity. The scRNA-seq dataset comprises 140 samples from 124 unique donors. For donors with multiple samples, we retained only the sample from the earliest timepoint. We then applied QUILT2 to sample-level BAM files across chromosome 6 using a reference panel constructed using whole genome sequencing samples from the 1000 Genomes Project phase 3 [16]. After imputation, we applied quality control to the single-nucleotide variants (SNVs). We retained variants with high imputation quality (INFO_SCORE > 0.8), excluded rare variants (minor allele frequency < 5% estimated with QUILT2 imputation results) and excluded variants significantly out of Hardy-Weinberg equilibrium (p < 1×10^-6^), yielding 76,833 variants for downstream analyses (**Supplementary Table 1**).

We next evaluated scTAPAS-derived genotype dosages against matched Global Screening Array (GSA) [5] and TOPMed-imputed genotypes available for the same donors [5], [17], [18], [19]. Across all scTAPAS-inferred variants, imputation quality scores (INFO_SCORE) from QUILT2 were positively correlated with genotype dosage concordance for both directly genotyped GSA (**Fig. 2a**; R^2^ = 0.702) and TOPMed-imputed variants (**Fig. 2b**; R^2^ = 0.765), supporting the use of this measure for variant filtering. We compared a less stringent filter of INFO_SCORE > 0.5 and a more stringent filter of INFO_SCORE > 0.8 with unfiltered variants. The INFO_SCORE > 0.8 filter led to strong dosage concordance with reference datasets, with median Pearson R^2^ of 0.770 for GSA variants and 0.771 for TOPMed-imputed variants (**Supplementary Table 2 & 3**), and substantially outperformed less stringent or no filtering (**Fig. 2c, d**). These comparisons support QUILT2 INFO_SCORE > 0.8 as a threshold for selecting scTAPAS-derived genotypes for downstream association analyses.

**Fig. 2.**
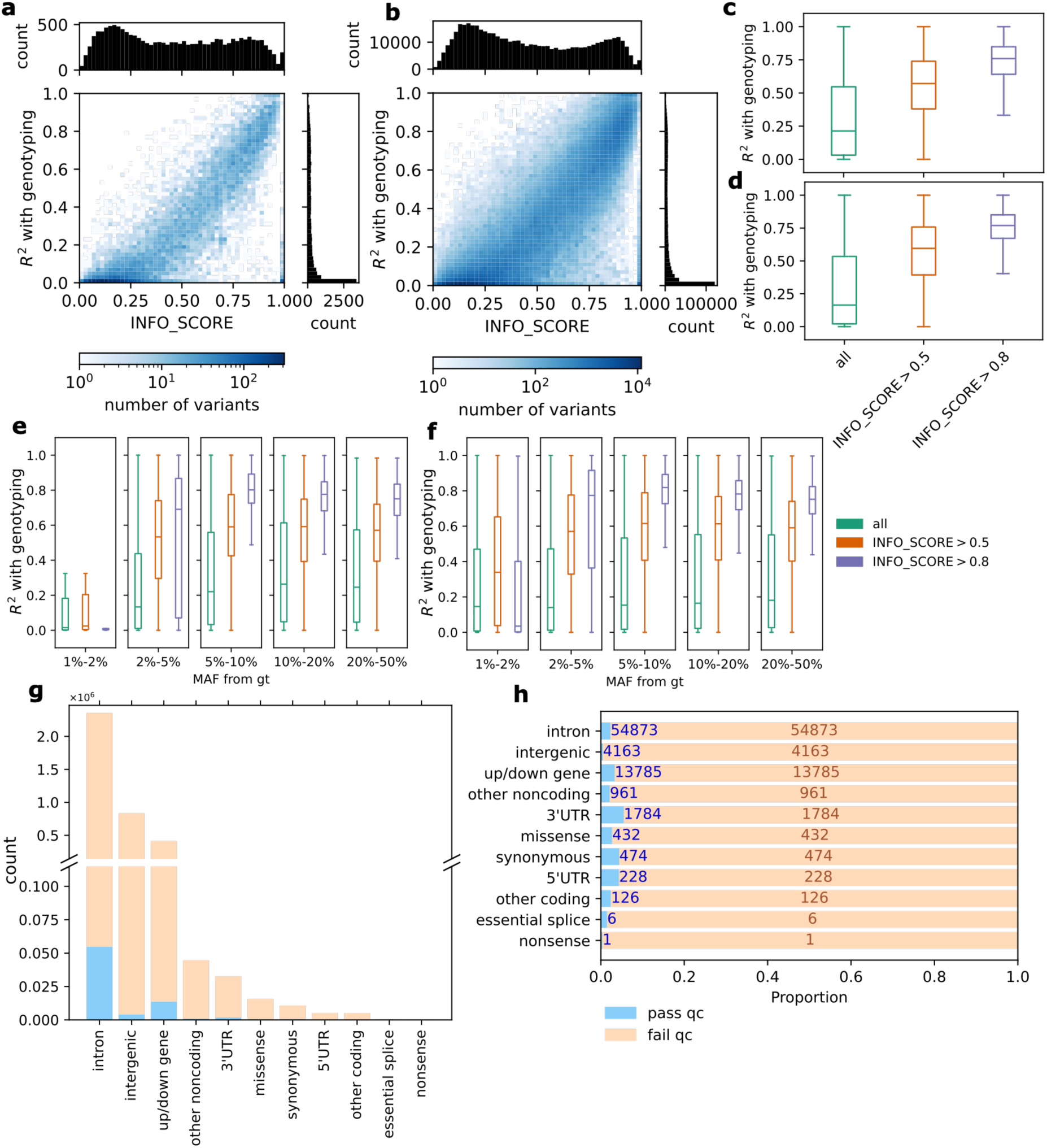
Evaluation and quality control of scTAPAS-inferred genotypes. Distribution of variants across QUILT2 quality scores (INFO_SCORE) and genotype dosage concordance between scTAPAS-inferred and matched reference genotypes for Global Screening Array (GSA) variants (**a**) and TOPMed-imputed variants (**b**). Marginal histograms show the distribution of INFO_SCORE and dosage concordance. Effect of INFO_SCORE filtering thresholds on genotype dosage concordance for GSA variants (**c**) and TOPMed-imputed variants (**d**). Genotype dosage concordance across minor allele frequency (MAF) bins and INFO_SCORE filtering thresholds for GSA variants (**e**) and TOPMed-imputed variants (**f**). Boxes show the median and interquartile range (IQR); whiskers extend to the most extreme values within 1.5 × IQR. Number (**g**) and proportion (**h**) of variants with each annotation category that pass or fail quality control.

In addition to INFO_SCORE, allele frequency was also a major indicator of imputation accuracy. scTAPAS inferred rare SNVs less accurately compared to common SNVs. We observed that SNVs with MAF > 5% in conventional genotyping showed substantially stronger concordance than those with MAF < 5% in combination with INFO_SCORE > 0.8 filtering (**Fig. 2e, f**). This pattern was consistent when allele frequency was estimated using genotyping array samples from the 1000 Genomes Project phase 3; SNVs with MAF > 5% and INFO_SCORE > 0.8 showed strong agreement with genotyping for GSA and TOPMed variants (**Fig. S1a, b**). We therefore excluded low-frequency SNVs from downstream analysis, using MAF > 5% in the scTAPAS-imputed variants as a conservative threshold for downstream analyses. This filter strengthened the association between INFO_SCORE and genotype dosage concordance for both GSA (R^2^=0.801) and TOPMed-imputed variants (R^2^=0.828; **Supplementary Table 4**). In addition, we confirmed that variants significantly out of Hardy-Weinberg equilibrium at p < 1×10^-6^ were poorly-imputed compared to other variants as expected (**Fig. S1c, d**). We therefore used INFO_SCORE > 0.8, MAF > 0.05 and HWE p > 1×10^-6^ as our default variant quality threshold to balance accuracy with SNV retention.

We next sought to understand how QUILT2 imputation quality is affected by single-cell sample quality, specifically the number of cells per sample. As expected, the logarithm of number of cells per sample was strongly positively correlated with sample-wise concordance with genotyping (R^2^ = 0.696, p = 1.27×10^-27^). Two samples with fewer than 500 cells, which were excluded from analysis in the original COMBAT study [14], showed particularly poor concordance and were excluded from downstream analyses (**Fig. S1e**). Among the remaining samples, number of cells remained positively, though less strongly, correlated with sample-wise concordance (R^2^ = 0.165, p = 2.78×10^-5^). This relationship likely reflects the increased read depth and coverage conferred by a greater number of cells when scRNA-seq reads are aggregated at the sample level. After exclusion of these two samples, scTAPAS variants were more accurately imputed overall: median Pearson R^2^ with array genotyping increased to 0.797 for GSA variants and 0.799 for TOPMed-imputed variants.

Finally, given the variable transcription of genomic regions and its potential impact of variable transcript expression on QUILT2 performance, we examined whether genomic annotation and transcript expression influenced imputation quality. We noted that most retained scTAPAS SNVs were annotated as intronic reflecting the overall abundance of intronic variants among RNA-derived sites (**Fig. 2g**), with higher proportions of other variant types such as 3’UTR variants passing quality control (**Fig. 2h**). Next, we asked whether this variation in passing rate was associated with transcription in profiled material. In COMBAT, which was generated from blood-derived scRNA-seq, annotation of SNVs to genes expressed in blood is strongly associated with passing quality control (β = 2.57, SE = 0.016), though being annotated to a gene not expressed in blood is also more weakly associated in comparison to intergenic SNVs (β = 0.972, SE = 0.017) (**Fig. S1f, g**). Together, these results indicate that successful genotype reconstruction from scRNA-seq depends on both the amount of sequencing information available per donor and the transcriptional context of individual variants.

### scTAPAS-derived genotypes support cell type-resolved eQTL discovery

We next tested whether scTAPAS-derived genotypes were sufficiently accurate to support cell type-resolved genetic association analysis. We benchmarked scTAPAS against the conventional array-based genotyping-plus-imputation strategy used in the COMBAT cohort [5]. Using 76,833 chromosome 6 variants that passed quality control (**Fig. 3a**) and 100 donors with more than 500 cells and matched genotype data, we performed *cis*-eQTL mapping using pseudobulk expression profiles across six major cell types, following the framework described in a previous COMBAT eQTL study [5]. Although we restricted eQTL analysis to chromosome 6 variants as a proof of principle, the scTAPAS framework can be applied genome-wide. Using the same modelling strategy, we fitted linear regression models for genotype-expression associations while adjusting for age, sex, the top five expression principal components (PCs) and the top two genetic PCs derived from scTAPAS genotypes.

**Fig. 3.**
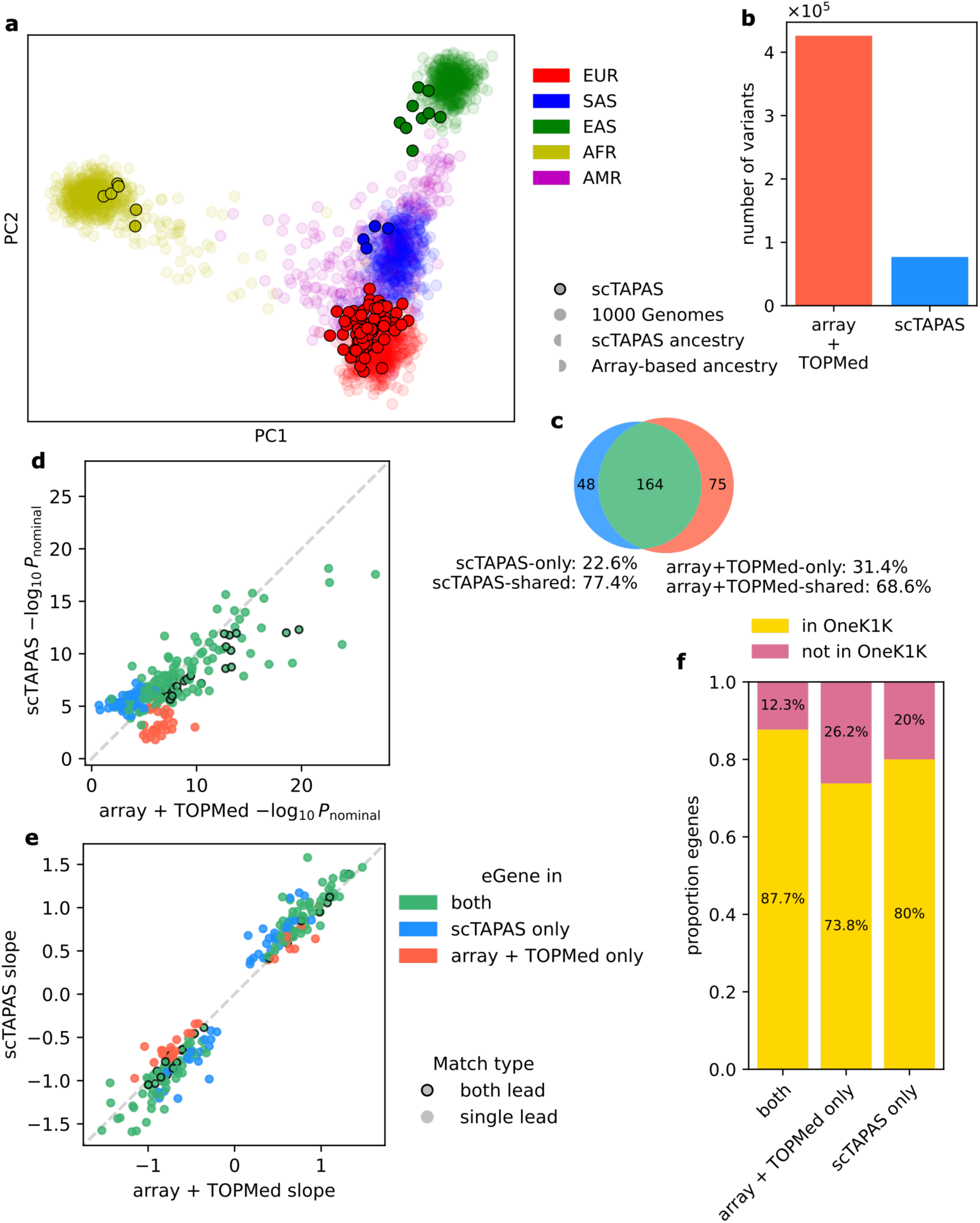
scTAPAS-derived genotypes recover cell type-resolved eQTL signals and donor genetic ancestry. Principal components 1 and 2 obtained using post-QC scTAPAS-derived variants (**a**). Each point represents one donor and is coloured by scTAPAS-derived and genotyping array-based ancestry assignments [5]. Number of variants tested in scTAPAS and array-based sc-eQTL analyses (**b**). Number of cell type-eGenes identified using scTAPAS-derived genotypes, array-based genotypes, or both (**c**). Percentage indicates the proportion exclusive to or shared between the two approaches. Nominal eQTL significance (-*log* p_nominal_) (**d**) and effect size (**e**) compared between scTAPAS and array-based analyses for shared and approach-specific cell type-eGene pairs. Black outlines indicate cell type-eGenes for which the same lead variant was identified in both analyses. Proportion of eGenes that were also supported by OneK1K or unique to each analysis (**f**).

Because population structure is an important potential confounder in genetic association studies, we first assessed whether it is possible to infer the major axes of donor genetic structure based on scTAPAS-derived genotypes. We calculated genetic PCs using 1000 Genomes Project reference samples and projected COMBAT donors into the same genetic space (**Fig. 3a**). Population assignments based on the distance of each donor to 1000 Genomes Project centroids of continental populations showed 100% concordance compared to non-outlier ancestry assignments previously inferred using matched array-based genotypes [5]. Thus, scTAPAS-derived genotypes can also provide reliable genetic PCs that can be used to adjust for ancestry-related structure in cohorts lacking conventional genotyping.

We then compared *cis*-eQTL discovery using scTAPAS-derived and conventional genotypes. scTAPAS provided fewer testable variants than array-based imputation, retaining 18.0% of the chromosome 6 variants used in the conventional COMBAT eQTL analysis (**Fig. 3b; Supplementary Table 6**). Despite this reduced variant coverage, scTAPAS recovered 68.6% of the cell type-eGene pairs, defined as genes with a significant variant association at 5% false discovery rate, using array-based imputed genotypes across the six cell types (**Fig. 3c, Supplementary Table 5**). We define cell type-eGene pairs as eGene discoveries matched by both gene and cell type, such that a gene detected in multiple cell types is counted separately in each cellular context. Of all the cell type-eGene pairs identified by scTAPAS and/or array genotyping-based sc-eQTLs, the same lead SNV was identified in 10.8% of cell type-eGene pairs. For 63.1% of cell type-eGene pairs, the lead SNV was tested in both modalities, whereas the remaining 26.1% could not be compared directly due to missing association statistics in one modality. Among the 164 cell type–eGene pairs identified by both approaches, we observed strong agreement in their nominal p values (Pearson R^2^ = 0.617, **Fig. 3d**) and effect sizes (Pearson R^2^ = 0.936, **Fig. 3e**) when the lead scTAPAS SNV was also tested in the array-based analysis.

We next examined cell type-eGene pairs detected by only one of the two approaches. The array-based imputation approach identified an additional 75 cell type-eGene pairs that were not detected with scTAPAS, which was expected due to the denser variant coverage and greater number of testable *cis*-variants available in the array-based imputation. Indeed, the array lead SNV was also tested in the scTAPAS analysis for only 29 (38.7%) of these cell type-eGene pairs, where effect sizes remained strongly correlated between the two approaches (Pearson R^2^ = 0.976, p = 2.29×10^-23^) despite differences in nominal p values (Pearson R^2^ = 0.0194, p = 0.472; **Fig. 3d**,**e**). These results suggest that many array-only signals reflect differences in variant coverage, with a smaller contribution from low-confidence variant calling. Conversely, scTAPAS identified 48 cell type-eGene pairs that did not reach significance in array-based analysis (**Fig. 3c**). For these scTAPAS-only eGenes where lead SNV was tested in both modalities, effect sizes remained strongly correlated between two approaches (Pearson R^2^ = 0.952, p = 2.47×10^-29^; **Fig. 3e**), whereas nominal p values showed only a weak positive correlation (Pearson R^2^ = 0.104, p = 0.0325; **Fig. 3d**). This suggests that differences in statistical significance may partly reflect differences in multiple-testing burden, although noise in some scTAPAS-derived variants may also contribute.

To further assess these signals, we compared scTAPAS-only eGenes with OneK1K, a large independent single-cell eQTL study of peripheral blood immune cells [2]. We found that 80.0% of the scTAPAS-only eGenes were also identified in OneK1K, compared with 73.8% of array-only eGenes (**Fig. 3f**). When requiring support in at least one matching cell type, we found 51.4% of scTAPAS-only eGenes and 52.4% of array-only eGenes were supported by OneK1K (**Fig. S2**). Together, these comparisons indicate that quality-controlled scTAPAS genotypes support reproducible cell type-resolved eQTL discovery despite substantially reduced variant density relative to conventional genotyping.

### scTAPAS enables HLA inference and HLA-TCR analysis

We next assessed whether scTAPAS-derived variants could support HLA imputation, having first verified adequate coverage of the classical HLA genes in sample-wise aggregated transcriptomic reads (**Fig. S4a-h**). Using scTAPAS-derived variants within the MHC region (chr6: 28,000,000 – 34,000,000), we applied EAGLE2 phasing [20] followed by HLA imputation using a multi-ancestry reference panel hosted on the Michigan Imputation Server (multi-ethnic HLA v2 (2020), [18], [19], [21]). The resulting HLA alleles were evaluated against matched HLA sequencing as the gold standard.

Across eight classical HLA genes, scTAPAS-derived and gold standard HLA sequencing dosage at two-field resolution showed modest correlation, with a median allele dosage correlation of R^2^ = 0.586 (95% quantile interval = (0, 1), **Fig. 4a, b**). This agreement improved when restricting the analysis to common HLA alleles with allele frequency over 5% (median R^2^ = 0.664; 95% quantile interval = (0.00,0.978)). scTAPAS outperforms a null model which assigns the most common genotype for each gene based on the gold standard, suggesting that scTAPAS provides information beyond population allele frequency alone (**Fig. 4a-b**). We also tested whether restricting the analysis to variants expressed in blood according to GTEx or COMBAT pseudobulks or to variants that are included in various genotyping arrays, could further improve HLA inference, but observed no further improvement (**Fig. S3c-d**).

**Fig. 4.**
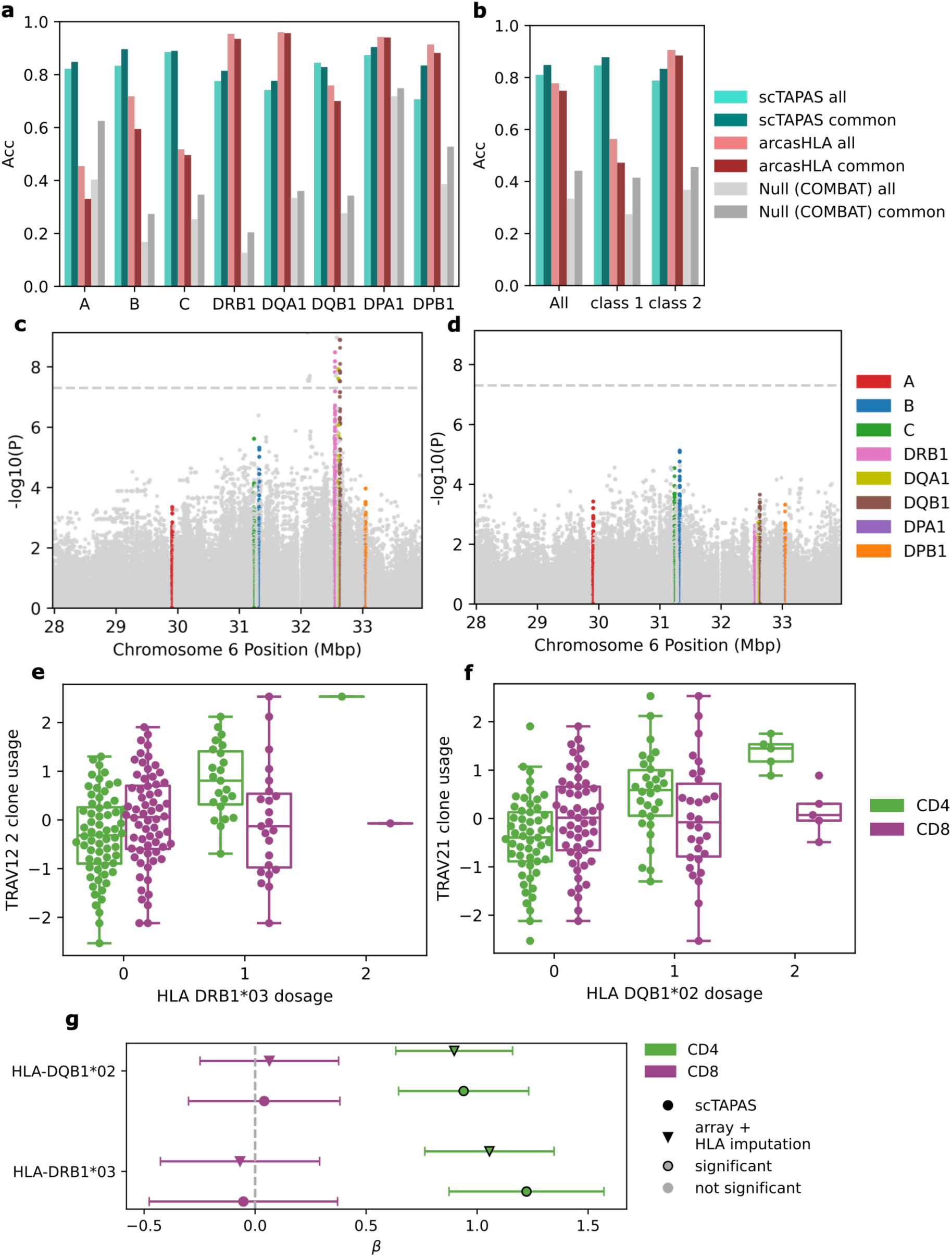
scTAPAS enables HLA imputation and HLA-TCR association analysis. HLA gene-wise accuracy for each classical HLA gene (**a**) and accuracy summarised across all classical HLA genes, class 1 and class 2 (**b**) inferred using scTAPAS, arcasHLA and null model at two-field resolution. Results are shown for all HLA alleles and for common alleles with allele frequency over 5% in the gold standard HLA sequencing. Nominal p-values from generalised linear model across MHC region variants combined for all TCR V gene segment usage in CD4+ T cells (**c**) and CD8+ T cells (**d**). HLA-TCR V gene associations for CD4+ T cells and CD8+ T cells, showing TRAV12-2 with HLA-DRB1*03 (**e**) and TRAV21 with HLA-DQB1*02 (**f**). Boxes show the median and interquartile range (IQR) where whiskers extend to the most extreme values within 1.5 × IQR, and all values are represented by scattered points. Effect sizes and 95% confidence intervals for HLA-TCR V gene associations identified using either scTAPAS- or array-based HLA imputation in CD4+ T cells, with the corresponding associations in CD8+ T cells (**g**).

To benchmark scTAPAS against an established approach for HLA inference from transcriptomic data, we compared scTAPAS-derived HLA alleles to those inferred by arcasHLA [22], a widely used method that directly infers HLA genotypes from RNA-seq reads and has previously been evaluated for application to scRNA-seq data [23]. Across eight classical HLA genes, we observed striking differences in relative performance between HLA class I (HLA-A, HLA-B and HLA-C) and class II (HLA-DQA1, HLA-DQB1, HLA-DRB1, HLA-DPA1, HLA-DPB1) loci. arcasHLA (**Supplementary Table 2**) performed better for class II genes (median R^2^ = 1.00 vs 0.481, for arcasHLA and scTAPAS, respectively), whereas scTAPAS showed higher accuracy for class I genes at two-field resolution (median R^2^ = 0.317 vs 0.659, for arcasHLA and scTAPAS, respectively, **Fig. 4b**). However, the two methods achieve similar results at one-field resolution (**Fig. S3a-b, Supplementary Table 7**). Reduced class I performance of arcasHLA in 5′ scRNA-seq has previously been reported [23].

Having established that scTAPAS could recover classical HLA variation, we next asked whether this information was sufficiently accurate for downstream immunogenetic association analysis. We tested association between imputed variants in the MHC region and TCR V gene segment usage using single-cell TCR sequencing data from COMBAT. We tested HLA-TCR association separately within conventional CD4+ and CD8+ T-cell subsets, using donor-level, clone-based TCR-V-gene usage and adjusting for age and sex (**Supplementary Table 8**). For CD4+ T cells, 43 TRAV and 44 TRBV gene segments were included in TCR-V-gene usage statistics. For CD8+ T cells, 42 TRAV and 44 TRBV gene segments were included. We observed significant associations between class II HLA variation and TCR V gene segment usage in CD4+ T cells (**Fig. 4c**), whereas we did not detect comparable associations between class I variation and TCR V gene segment usage in CD8+ T cells (**Fig. 4d**). The absence of significant CD8+ T-cell associations may in part reflect the larger number of CD4+ than CD8+ T cells available in COMBAT (ratio = 2.56:1) [14].

Among the strongest CD4+ T-cell associations, HLA-DRB1*03 was associated with TRAV12-2 usage (*β* = 1.22, p = 1.05×10^-9^) in CD4+ T cells but not CD8+ T cells (*β* = -0.0527 p = 0.808, **Fig. 4e**). Additionally, HLA-DQB1*02 was associated with TRAV21 usage (*β* = 0.940, p = 1.42×10^-8^) in CD4+ T cells but not CD8+ T cells (*β* = 0.0410, p = 0.814, **Fig. 4f**). These results are consistent with an equivalent analysis based on HLA imputation of GSA variants for the same samples, both in the significant HLA-TCR V gene pairs as well as effect size and significance (**Fig. 4g; Supplementary Table 9**). Together, these results show that scTAPAS can support HLA-aware immunogenetic analyses of matched single-cell TCR sequencing data.

## Discussion

Our results show that genetic information contained within scRNA-seq reads can be used to reconstruct genotypes at the donor level, enabling downstream genetic and immunogenetic association analyses. Using matched scRNA-seq and genotyping data from COMBAT, we demonstrate that scTAPAS variant dosage showed sufficient concordance with array-based genotyping. We rediscover 68.6% of cell type-eGenes in COMBAT *cis*-eQTLs despite only testing 18.0% of variants compared to genotyping, indicating that scTAPAS genotyping is sufficient to capture a substantial proportion of the *cis*-regulatory architecture of the genome. We further extended this approach to the MHC region, showing that scRNA-seq-derived genetic variation can support classical HLA imputation, enable HLA-TCR association analysis and may complement existing approaches to RNA-based HLA imputation. Together, we show that scRNA-seq is a potential source of donor genotyping for single-cell cohorts in which conventional genotyping or HLA typing is unavailable.

scTAPAS could be applied to scRNA-seq cohorts lacking matched genotyping data, provided the necessary consent and governance for genetic analysis are in place. Augmenting these datasets may be particularly valuable for genetic and immunogenetic analyses that combine cohorts from multiple sources, potentially increasing the number and size of datasets available for analysis. We have also shown that scTAPAS-derived variants can be used to infer donor genetic ancestry, providing covariates that may be important for controlling for population structure in statistical models. The ability to infer HLA variation provides a distinct extension of this framework beyond conventional eQTL analysis.

HLA type is a crucial determinant of the TCR repertoire, as HLA and TCR molecules interact through antigen presentation and recognition during both T-cell development and immune responses. This information is particularly relevant to single-cell TCR-seq studies, which increasingly provide paired measurements of TCR sequences and transcriptional cell state but do not necessarily include conventional HLA typing or genotyping [14], [24], [25], [26]. scTAPAS therefore provides a practical alternative for HLA inference where HLA typing or genotyping is unavailable due to cost or technical reasons. In COMBAT, HLA variation inferred using scTAPAS reproduced CD4+ T-cell associations between HLA class II alleles and TCR V-gene usage observed using conventional genotype-derived HLA imputation. These findings illustrate how genetic information recovered from matched scRNA-seq can provide immunogenetic context for single-cell immune-repertoire studies.

The contrasting performance of scTAPAS and arcasHLA across classical HLA genes at two-field resolution further suggests that the two approaches capture complementary information from scRNA-seq. arcasHLA directly uses reads derived from HLA transcripts, whereas scTAPAS reconstructs variation across the surrounding MHC region and uses population haplotype structure to impute HLA alleles. Consistent with previous benchmarking in 5’ scRNA-seq, we observed lower two-field performance of arcasHLA for class I than class II loci [23], despite strong class I performance of arcasHLA using well-covered bulk RNA-seq [22], [27]. Given that two-field HLA resolution can be crucial for applications requiring fine-grained HLA stratification, including stratification by autoantibody risk [28], we suggest that scTAPAS may complement direct transcript-based approaches such as arcasHLA for HLA inference from scRNA-seq.

Several limitations should be considered when applying scTAPAS to other scRNA-seq datasets. First, we applied and evaluated scTAPAS using the COMBAT dataset, specifically the 10x Genomics Chromium (v1.1) 5’ scRNA-seq from peripheral blood mononuclear cells, and performance may differ across sequencing chemistries and biological tissues. Because genotype inference depends on genetic information captured within sequencing reads, differences in read length, transcript coverage and gene expression are likely to influence which variants can be accurately inferred. Whether comparable performance can be achieved using 3′ scRNA-seq therefore requires direct evaluation. Second, genotype concordance increased with the number of cells available per donor, though to a lesser extent above a threshold of 500 cells, indicating that datasets with low cell numbers or sequencing depth may provide insufficient genetic information for reliable imputation. Third, genotype inference was less accurate for rare variants, and our downstream analyses consequently focused on common variants. scTAPAS is therefore unlikely to provide the coverage required for analyses centered on rare genetic variation. Fourth, our downstream genetic association analyses were restricted to chromosome 6. Although the same genotype-reconstruction framework can in principle be applied across all autosomes, its performance and computational requirements at genome-wide scale remain to be systematically evaluated. Fifth, scTAPAS takes reads aligned to a single linear reference genome as input, which may introduce haplotype-dependent alignment bias in regions of structural variation, such as the HLA-DRB genes, and may account for the higher HLA-DRB1 accuracy of arcasHLA relative to scTAPAS. Finally, reference-based imputation depends on representation of donor haplotypes within the reference panel, and performance may vary across ancestries that are less well represented in available reference resources. Evaluation in larger and more ancestrally diverse single-cell cohorts will therefore be important for defining the generalisability of the approach.

Despite these limitations, scTAPAS provides a route to extend genetic analysis to existing and future scRNA-seq datasets for which matched genotype data are unavailable. In addition to enabling cell type-resolved eQTL mapping, reconstructed genotypes can provide genetic ancestry information and support HLA-aware analyses of matched immune-repertoire data. This may be particularly valuable when conventional genotyping or HLA typing is constrained by cost, limited availability of biological material or the design of existing studies, and when integrating single-cell datasets across cohorts. Because genotype reconstruction from transcriptomic data generates potentially identifiable genetic information, its application should be restricted to datasets with appropriate participant consent and governance for genetic analysis. Overall, scTAPAS provides a practical workflow for leveraging scRNA-seq to study genetic regulation and adds immunogenetic context to matched single-cell immune repertoire data.

## Acknowledgements

This work was funded by a Kennedy Trust Senior Research Fellowship award (KENN202109). LJD and SS are supported by the Kennedy Trust for Rheumatology (KENN71803). This project was supported by the University of Oxford COVID-19 Research Response Fund and NIHR Oxford Biomedical Research Centre. This project has been funded in whole or in part with Federal funds from the Center for Cancer Research, National Cancer Institute, National Institutes of Health, Department of Health and Human Services, under Contract No. 75N91019D00024. The content of this publication does not necessarily reflect the views or policies of the Department of Health and Human Services, nor does mention of trade names, commercial products, or organizations imply endorsement by the U.S. Government.

## Author Contributions

Conceptualisation: YL, HK; Data Curation: MNC, PJN, KMK, AYC, AJM, RAW, BPF, SNS, CAD, COMBAT consortium; Formal analysis: HK, JA; Methodology: HK, YL, JA, LJD, AJM, SNS, GB, ZZ, ESN; Software: HK; Supervision: YL, LJD; Visualisation: HK, YL; Writing-original draft preparation: HK, YL; all authors contributed to the review and editing of the manuscript.

## Code Availability

scTAPAS is available via GitHub (https://github.com/yang-luo-lab/scTAPAS/tree/main), included with scripts used to generate manuscript figures.

## Data Availability

COMBAT 5’ single-cell RNA sequencing (EGA: EGAD00001008007), matched array-based genotyping (EGA: TBC) and single-cell TCR sequencing (EGA: EGAD00001007965) are available. HLA sequencing will be available upon publication of manuscript.

High-coverage whole genome sequencing [16] and genotyping array data from 1000 Genomes Project phase 3 are publicly available [29].

## Methods

### Data collection and quality control

We applied scTAPAS to data from the COVID-19 Multi-omic Blood Atlas (COMBAT) [5], [14], including 5’ single-cell RNA sequencing (EGA: EGAD00001008007), matched array-based genotyping (EGA: TBC), HLA typing and single-cell TCR sequencing (EGA: EGAD00001007965). We used high-coverage whole-genome sequencing from the 1000 Genomes Project v3 [16] as a reference panel for genotype imputation with QUILT2.

### Single-cell RNA sequencing

COMBAT peripheral blood mononuclear cells (PBMCs) were profiled using the 10x Genomics Chromium (v1.1) 5’ [14]. Reads from scRNA-seq data were previously processed using CellRanger (v3.1.0) and stored as *.bam* files [14]. To refine cell-donor assignments, we applied the Souporcell3 demultiplexing pipeline with `skip-remap` and `common-variants` options [11], [30], specifying variants derived from the post-quality control set of TOPMed-imputed GSA variants, with any monomorphic variants excluded [5], [14]. We next matched Souporcell cluster genotypes to sample identifiers where possible using Pearson correlation with array-based genotypes of samples known to be included in each pool, and extracted chromosome 6 reads in sample-wise *.bam* format based on the results [5], [14], [31].

### Array genotyping

For scTAPAS evaluation, we utilise previously generated genotyping array data from COMBAT [5]. Genotyping was performed using the Global Screening Array-24 v3 (Illumina, San Diego, CA, USA). Quality control was applied for variants and samples, including removal of sex mismatched samples, samples with low call rate (<98%), samples with outlying heterozygosity, variants with low call rate (<98%), rare variants (<2% MAF), variants out of Hardy-Weinberg equilibrium (p<0.001), indels, copy number variation and strand-ambiguous SNVs [5]. TOPMed imputation was performed, and additional filtering on R^2^ and MAF was applied [5]. We used directly genotyped GSA variants and TOPMed-imputed variants to assess concordance with scTAPAS-derived genotype dosages.

### HLA sequencing

For evaluation of scTAPAS HLA inference, we used HLA sequencing as the gold standard. HLA sequencing was performed independently using two sequencing methods, including clinical grade targeted next-generation sequencing (NGS) and biotinylated DNA probe enrichment [32], to ensure a high quality call-set. Where HLA types were inconsistent between these, we used the clinical grade calls.

Clinical grade HLA typing was performed using a targeted NGS method. Briefly, locus-specific primers are used to amplify a total of 26 polymorphic exons of HLA-A and B (exons 1 to 4), HLA-C (exons 1 to 5), HLA-E (exon 3), DPA1 (exon 2), DPB1 (exons 2 to 4), DQA1 (exon 1 to 3), DQB1 (exons 2 & 3), DRB1 (exons 2 and 3), and DRB3/4/5 (exon 2) genes with Fluidigm Access Array System with Juno LP 48.48 IFCs (Standard Bio Tools, Inc., South San Francisco, CA 94080 USA). The 26 Fluidigm PCR amplicons are pooled and subjected to sequencing on an Illumina MiSeq sequencer (Illumina, San Diego, CA 92122 USA). HLA alleles and genotypes are called using the Omixon HLA Explore (version 2.0.0) software (Omixon, Budapest, Hungary).

Additionally, an established biotinylated DNA probe enrichment method for sequencing HLA genes was used [32]. To generate sequencing libraries, the following modifications were made: 500ng of genomic DNA was fragmented using NEBNext® Ultra II FS module (E7810, New England Biolabs, Ipswich, MA) and ligated with IDT xGen Stubby Adapters (Integrated DNA Technologies, Coralville, IA) by NEBNext® Ultra II Ligation module (E7959, New England Biolabs, Ipswich, MA). Dual size selection with Agencourt AMPure XP beads (Beckman Coulter Life Sciences, Indianapolis, IN) was used to select 900 bp DNA fragments. Samples were molecularly tagged with 5ul of 10uM unique dual-index primers (Integrated DNA Technologies, Coralville, IA). Then, post-amplification was performed based on the Kapa HyperPrep protocol (Kapa Biosystems, Wilmington, MA). Equivalent quantities of each DNA library were pooled for enrichment, which was performed using the capture probes and a modified version of the Nextera Rapid Capture enrichment protocol (Illumina) as described [32]. Sequencing was performed using the NovaSeq 6000 platform with an SP flowcell (Illumina Inc., San Diego, CA) with 2×250bp reads. HLA class I and II alleles were determined using NGSengine 2.30.0 (GenDX, Utrecht, Netherlands).

### Single-cell TCR sequencing

Single-cell TCRs were previously profiled using 10x Genomics Chromium (v1.1) for the same cells as for COMBAT scRNA-seq, and processed using CellRanger VDJ [14]. For downstream analysis, we retained only T cells with TCRs with a single alpha chain and single beta chain. Of those remaining, we observed T cells with identical TCRs on the nucleotide level across different samples but originating from the same pool. These likely occur due to incorrect assignment of cells to individuals rather than true TCR sharing. We assigned these inter-sample single-pool T-cell clones to the sample with the greatest number of members, and removed other member cells that were assigned to other samples. Where there was a tie in the greatest number, all cells from the inter-sample single-pool T-cell clone were excluded.

### scTAPAS pipelines

The majority of scTAPAS was implemented using Nextflow (version 24.04.2) [33], inclusive of reference panel preparation, genotype inference with QUILT2, ancestry inference, sc-eQTL analysis, and HLA-TCR association analysis.

### Preparing a reference panel for QUILT2 imputation

We used high-coverage whole-genome sequencing samples from 1000 Genomes Project phase 3 [16] to prepare a reference panel for QUILT2 imputation. We removed samples NA12878, NA12891 and NA12892, split multiallelic records and normalised variants to a reference fasta using BCFtools (version 1.18) [31]. After calculating chromosome 6 chunks using `QUILT::quilt_chunk_map`, we prepared a QUILT2 reference for each of the overlapping chunks using `QUILT2_prepare_reference` with the processed 1000 Genomes Project panel.

### Inferring donor genotypes using scTAPAS

We applied QUILT2 (version 2.02) [13] to sample-wise BAM files containing chromosome 6-aligned scRNA-seq reads from COMBAT. QUILT2 was run with settings `method = diploid`, `nGen = 100` and `buffer = 500000` for 140 scRNA-seq samples in COMBAT. We specified the genetic map file as the genetic recombination map file included in the QUILT2 package which is specific for CEU population. 35 QUILT2 jobs, one for each chromosome 6 chunk, were run on 8 cores with 160 GB of memory, with one job failing and retrying with 184 GB. On average each job took 3 hours, 53 minutes and 18 seconds. We selected 102 samples from unique donors with matches in the genotyping array, taking the sample from the first timepoint if a donor had given multiple samples. With the variants imputed by QUILT2, we assembled VCF files for chromosome 6, first removing overlapping regions of each chunk and concatenating with BCFtools (version 1.18) [31]. We applied filters using BCFtools view, keeping variants with internal QUILT2 imputation quality measure INFO_SCORE > 0.8, minor allele frequency MAF > 0.05 and Hardy-Weinberg equilibrium p value > 1×10^-6^.

### Comparison of scTAPAS genotypes to array-based genotypes

We evaluated SNVs from scTAPAS prior to quality control through comparison to array-based genotypes. We calculated Pearson’s correlation coefficient for each variant between the scTAPAS-derived and genotype dosage and either directly genotyped GSA or TOPMed-imputed genotype dosage across donors. For each variant, we excluded samples with missing scTAPAS or genotype dosage. For variants that are monomorphic in scTAPAS or TOPMed-imputed genotyping, we estimated Pearson’s correlation coefficient as zero.

We contextualised the imputation quality of the SNVs by obtaining allele frequencies from COMBAT genotyping [5] and 1000 Genomes Project phase 3 [16], as well as variant annotations from Ensembl VEP [34], gene expression data from GTEx [35], [36] and gene expression data from COMBAT scRNA-seq pseudobulks [14]. We applied Ensembl VEP to all pre-QC scTAPAS SNVs with options `pick`, `check_existing`, `species = homo_sapiens`, `assembly = GRCh38`. We simplified annotations to categories essential splice, nonsense, missense, synonymous, other coding, 5’ UTR, 3’UTR, intron, other noncoding, upstream/downstream of gene and intergenic (**Supplementary Table 10**). For variants labelled with more than one annotation, we took the one that appeared first in the previous list of simplified annotations. For gene annotations from Ensembl VEP we labelled those that were expressed in blood if median expression was one or more across samples. SNVs were otherwise labelled with a gene not expressed in blood, or as intergenic. Finally, we fitted a logistic regression model for all SNVs as below to test association of variant gene expression status with the SNV passing quality control.

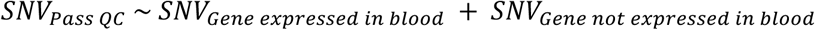

### scTAPAS ancestry assignment

We downloaded phase 3 1000 Genomes Project array genotyping samples in Plink2 binary format [29]. After extracting unrelated samples, we calculated the intersection of variants with scTAPAS, excluding the MHC region. Using Plink 2 alpha 3.7 [37], we fitted a generalised linear model with 1000 Genomes Project or COMBAT respectively as phenotype to identify variants that differ significantly between the modalities with p < 1×10^-6^. We excluded these variants, and pruned for linkage disequilibrium using Plink settings of `window size = 50`, `shift = 5` and `VIF = 0.2` based on 1000 Genomes Project samples. Next, we calculated principal components based on these 1000 Genomes Project samples with pruned variants, and projected both 1000 Genomes Project and COMBAT scTAPAS samples onto them. Each of the first 10 principal components (PCs) were normalised by standard deviation from 1000 Genomes Project principal components. For each COMBAT sample, we calculated the sum of absolute distances across these 10 normalised PCs to each 1000 Genomes Project population centroid, with the minimum distance used to assign a sample to a population. We compared these assignments to those derived originally from COMBAT genotyping, excluding samples previously identified as ancestry outliers [5].

### scTAPAS sc-eQTL analysis

Using scTAPAS-derived genotypes, we conducted sc-eQTL analysis for six cell types using TensorQTL (version 1.0.10) [38], following the approach described previously for COMBAT [5]. We repeated the COMBAT analysis [5] using the updated TensorQTL version but restricted the analysis to chromosome 6 only. Briefly, gene expression profiles based on pseudobulk single-cell gene expression for each cell type were used as the phenotype in TensorQTL cis-mapping analysis. scTAPAS SNVs were used to calculate genetic principal components using Plink 2 alpha 3.7 [37]. For each genotype-expression association, we adjusted for age, sex, the top two genetic PCs and the top five expression PCs. We ran TensorQTL `cis_nominal` and `cis` mode with `qval_lambda = 0.85`, where false discovery rate was controlled at 5% [38], [39]. We finally ran `cis-independent` mode to identify independent cis-eQTLs.

For evaluation we compared cell type-eGene pairs identified in cis independent mode, recording which cell type-eGenes were identified with scTAPAS and genotype-based eQTL analysis. We compared nominal p values and slope from cis nominal mode per eGene and cell type, taking the lead SNV for any eGene identified in scTAPAS, and comparing to the same SNV in the genotyping-based results where they existed. For eGenes identified only in array-based genotyping, we took the lead variant from these sc-eQTLs, and compared with the equivalent scTAPAS variant where it was tested in scTAPAS sc-eQTLs. We also compared eGenes identified by each approach with those reported in OneK1K, recording both overall eGene support and support within a matching cell type [2].

### Inferring HLA alleles and MHC region SNVs from scRNA-seq

We imputed classical HLA alleles and variants across the MHC region using scTAPAS-derived SNVs. Prior to imputation, we extracted SNVs within the MHC region (chr6: 28,000,000 – 34,000,000) that were present on the HLA imputation panel. We performed HLA imputation using the multi-ancestry reference panel (multi-ethnic HLA v2 (2020)) [21] hosted on Michigan Imputation Server [18], [19] with EAGLE2 phasing [20]. To investigate whether variant selection influenced HLA imputation performance, we additionally restricted scTAPAS-derived variants to those included in the Infinium Global Screening Array-24 v3, Infinium Immuno Array-24 v2, Human Core Exome-12 v1 BeadChip (all Illumina, San Diego, CA, USA), as well as variants annotated to genes expressed in blood according to GTEx or COMBAT pseudobulked gene expression data, where median counts greater than 1 across samples.

We assessed the quality of imputation of HLA alleles against HLA sequencing as a gold standard. As benchmarks, we include results from arcasHLA [22] version 0.6.0 applied to per-sample *.bam* files as input to infer classical HLA genes. Additionally, we implemented a null model that predicts the most common genotype observed in the gold standard for each classical HLA allele. For scTAPAS, arcasHLA and the null model, we calculated Pearson correlation between dosage of each predicted allele and the gold standard allele. We also calculated classical HLA gene-wise accuracy, HLA class I and HLA class II accuracy, as well as overall HLA type accuracy, defined below.

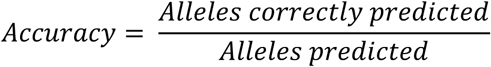

These calculations were performed for two-field resolution and one-field resolution HLA alleles. We also calculated HLA accuracy based only on common, with MAF > 5% in the gold standard at each resolution. We implemented this by excluding HLA alleles with MAF < 5% from the predicted and gold standard set. As such, in the unusual case where an individual is homozygous for a rare HLA allele, their sample would not be used to calculate the relevant HLA gene accuracy for common alleles. However, an individual heterozygous for the same rare allele would be included provided their other allele is common.

### HLA-TCR association testing for T-cell subsets

We used scTAPAS-derived variants within the MHC region and classical HLA alleles to model COMBAT TCR V gene usage using generalised linear model with age and sex as covariates, as below.

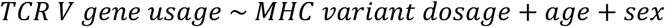

We calculated V gene usage per T-cell clone for samples in COMBAT [14]. T-cell clones were identified based on identical V gene, J gene and amino acid CDR3 of TCRs observed in T cells from the same individual. For CD4+ T cells and CD8+ T cells, we calculated V gene usage based on clones with at least one CD4+ or CD8+ T cell member respectively. TCR V gene usage was normalised to represent the proportion of clones using the V gene per individual, and further transformed using the inverse normal rank transformation. We tested only V genes with non-zero usage identified in at least 5% of samples, and excluded variants with MAF < 5%.

## References

[1] LifeLines Cohort Study et al., ‘Single-cell RNA sequencing identifies celltype-specific cis-eQTLs and co-expression QTLs’, Nat. Genet., vol. 50, no. 4, pp. 493–497, Apr. 2018, doi: 10.1038/s41588-018-0089-9.

[2] S. Yazar et al., ‘Single-cell eQTL mapping identifies cell type–specific genetic control of autoimmune disease’, Science, vol. 376, no. 6589, p. eabf3041, Apr. 2022, doi: 10.1126/science.abf3041.

[3] J. B. Kang et al., ‘Mapping the dynamic genetic regulatory architecture of HLA genes at single-cell resolution’, Nat. Genet., vol. 55, no. 12, pp. 2255–2268, Dec. 2023, doi: 10.1038/s41588-023-01586-6.

[4] D. J. Taylor et al., ‘Sources of gene expression variation in a globally diverse human cohort’, Nature, vol. 632, no. 8023, pp. 122–130, Aug. 2024, doi: 10.1038/s41586-024-07708-2.

[5] J. Astley et al., ‘Single-cell eQTL mapping identifies disease-specific genetic control of COVID-19 infection’, Jun. 30, 2025, medRxiv. doi: 10.1101/2025.06.30.25330463.

[6] A. S. E. Cuomo et al., ‘Impact of Rare and Common Genetic Variation on Cell Type-Specific Gene Expression’, Mar. 21, 2025. doi: 10.1101/2025.03.20.25324352.

[7] E. Garrison and G. Marth, ‘Haplotype-based variant detection from short-read sequencing’, Jul. 20, 2012, arXiv: arXiv:1207.3907. doi: 10.48550/arXiv.1207.3907.

[8] R. Poplin, et al., ‘Scaling accurate genetic variant discovery to tens of thousands of samples’, Jul. 24, 2018, bioRxiv. doi: 10.1101/201178.

[9] X. Huang and Y. Huang, ‘Cellsnp-lite: an efficient tool for genotyping single cells’, Bioinformatics, vol. 37, no. 23, pp. 4569–4571, Dec. 2021, doi: 10.1093/bioinformatics/btab358.

[10] F. Liu et al., ‘Systematic comparative analysis of single-nucleotide variant detection methods from single-cell RNA sequencing data’, Genome Biol., vol. 20, no. 1, p. 242, Nov. 2019, doi: 10.1186/s13059-019-1863-4.

[11] H. Heaton et al., ‘Souporcell: robust clustering of single-cell RNA-seq data by genotype without reference genotypes’, Nat. Methods, vol. 17, no. 6, pp. 615–620, Jun. 2020, doi: 10.1038/s41592-020-0820-1.

[12] S. A. Biagini et al., ‘Genotype imputation from low-coverage data for medical and population genetic analyses’, Genome Res., vol. 35, no. 9, pp. 1929–1941, Sep. 2025, doi: 10.1101/gr.280175.124.

[13] Z. Li, A. Albrechtsen, and R. W. Davies, ‘Flexible read-aware genotype imputation from sequence using biobank sized reference panels’, Nat. Commun., vol. 17, no. 1, p. 524, Dec. 2025, doi: 10.1038/s41467-025-67218-1.

[14] COvid-19 Multi-omics Blood ATlas (COMBAT) Consortium, ‘A blood atlas of COVID-19 defines hallmarks of disease severity and specificity’, Cell, vol. 185, no. 5, pp. 916–938.e58, Mar. 2022, doi: 10.1016/j.cell.2022.01.012.

[15] R. W. Davies et al., ‘Rapid genotype imputation from sequence with reference panels’, Nat. Genet., vol. 53, no. 7, pp. 1104–1111, Jul. 2021, doi: 10.1038/s41588-021-00877-0.

[16] M. Byrska-Bishop et al., ‘High-coverage whole-genome sequencing of the expanded 1000 Genomes Project cohort including 602 trios’, Cell, vol. 185, no. 18, pp. 3426–3440.e19, Sep. 2022, doi: 10.1016/j.cell.2022.08.004.

[17] D. Taliun et al., ‘Sequencing of 53,831 diverse genomes from the NHLBI TOPMed Program’, Nature, vol. 590, no. 7845, pp. 290–299, Feb. 2021, doi: 10.1038/s41586-021-03205-y.

[18] S. Das et al., ‘Next-generation genotype imputation service and methods’, Nat. Genet., vol. 48, no. 10, pp. 1284–1287, Oct. 2016, doi: 10.1038/ng.3656.

[19] C. Fuchsberger, G. R. Abecasis, and D. A. Hinds, ‘minimac2: faster genotype imputation’, Bioinformatics, vol. 31, no. 5, pp. 782–784, Mar. 2015, doi: 10.1093/bioinformatics/btu704.

[20] P.-R. Loh et al., ‘Reference-based phasing using the Haplotype Reference Consortium panel’, Nat. Genet., vol. 48, no. 11, pp. 1443–1448, Nov. 2016, doi: 10.1038/ng.3679.

[21] Y. Luo et al., ‘A high-resolution HLA reference panel capturing global population diversity enables multi-ancestry fine-mapping in HIV host response’, Nat. Genet., vol. 53, no. 10, Art. no. 10, Oct. 2021, doi: 10.1038/s41588-021-00935-7.

[22] R. Orenbuch, I. Filip, D. Comito, J. Shaman, I. Pe’er, and R. Rabadan, ‘arcasHLA: high-resolution HLA typing from RNAseq’, Bioinformatics, vol. 36, no. 1, pp. 33–40, Jan. 2020, doi: 10.1093/bioinformatics/btz474.

[23] B. D. Solomon et al., ‘Prediction of HLA genotypes from single-cell transcriptome data’, Front. Immunol., vol. 14, Apr. 2023, doi: 10.3389/fimmu.2023.1146826.

[24] G. Dunlap et al., ‘Clonal associations between lymphocyte subsets and functional states in rheumatoid arthritis synovium’, Nat. Commun., vol. 15, no. 1, p. 4991, Jun. 2024, doi: 10.1038/s41467-024-49186-0.

[25] F. Hayashi et al., ‘Antigen specificity of clonally enriched CD8+ T cells in multiple sclerosis’, Nat. Immunol., pp. 1–13, Feb. 2026, doi: 10.1038/s41590-025-02412-3.

[26] E. Stephenson et al., ‘Temporal multi-omics analysis of COVID-19 in end-stage kidney disease’, Cell Genomics, vol. 5, no. 8, Aug. 2025, doi: 10.1016/j.xgen.2025.100918.

[27] L. Song, G. Bai, X. S. Liu, B. Li, and H. Li, ‘Efficient and accurate KIR and HLA genotyping with massively parallel sequencing data’, Genome Res., vol. 33, no. 6, pp. 923–931, Jul. 2023, doi: 10.1101/gr.277585.122.

[28] N. Gharahdaghi et al., ‘Interleukin-10 Autoantibodies and HLA-DRB1*01:03 in Inflammatory Bowel Disease’, N. Engl. J. Med., vol. 394, no. 22, pp. 2212–2222, Jun. 2026, doi: 10.1056/NEJMoa2513654.

[29] A. Auton et al., ‘A global reference for human genetic variation’, Nature, vol. 526, no. 7571, pp. 68–74, Oct. 2015, doi: 10.1038/nature15393.

[30] M. Weerakoon, H. Vu, R. Behboudi, and H. Heaton, ‘Souporcell3: robust demultiplexing for high-donor single-cell RNA-seq datasets’, Bioinformatics, vol. 42, no. 3, p. btag117, Mar. 2026, doi: 10.1093/bioinformatics/btag117.

[31] P. Danecek et al., ‘Twelve years of SAMtools and BCFtools’, GigaScience, vol. 10, no. 2, p. giab008, Jan. 2021, doi: 10.1093/gigascience/giab008.

[32] P. J. Norman et al., ‘Defining KIR and HLA Class I Genotypes at Highest Resolution via High-Throughput Sequencing’, Am. J. Hum. Genet., vol. 99, no. 2, pp. 375–391, Aug. 2016, doi: 10.1016/j.ajhg.2016.06.023.

[33] P. Di Tommaso, M. Chatzou, E. W. Floden, P. P. Barja, E. Palumbo, and C. Notredame, ‘Nextflow enables reproducible computational workflows’, Nat. Biotechnol., vol. 35, no. 4, pp. 316–319, Apr. 2017, doi: 10.1038/nbt.3820.

[34] W. McLaren et al., ‘The Ensembl Variant Effect Predictor’, Genome Biol., vol. 17, no. 1, p. 122, Jun. 2016, doi: 10.1186/s13059-016-0974-4.

[35] THE GTEX CONSORTIUM, ‘The GTEx Consortium atlas of genetic regulatory effects across human tissues’, Science, vol. 369, no. 6509, pp. 1318–1330, Sep. 2020, doi: 10.1126/science.aaz1776.

[36] THE GTEX CONSORTIUM, ‘Genetic effects on gene expression across human tissues’, Nature, vol. 550, no. 7675, pp. 204–213, Oct. 2017, doi: 10.1038/nature24277.

[37] C. C. Chang, C. C. Chow, L. C. Tellier, S. Vattikuti, S. M. Purcell, and J. J. Lee, ‘Second-generation PLINK: rising to the challenge of larger and richer datasets’, GigaScience, vol. 4, no. 1, pp. s13742-015-0047–8, Dec. 2015, doi: 10.1186/s13742-015-0047-8.

[38] A. Taylor-Weiner et al., ‘Scaling computational genomics to millions of individuals with GPUs’, Genome Biol., vol. 20, no. 1, p. 228, Nov. 2019, doi: 10.1186/s13059-019-1836-7.

[39] H. Ongen, A. Buil, A. A. Brown, E. T. Dermitzakis, and O. Delaneau, ‘Fast and efficient QTL mapper for thousands of molecular phenotypes’, Bioinformatics, vol. 32, no. 10, pp. 1479–1485, May 2016, doi: 10.1093/bioinformatics/btv722.

