## supplementary figures for "Reconstructing donor genotypes from scRNA-seq for downstream eQTL and HLA–TCR analysis"

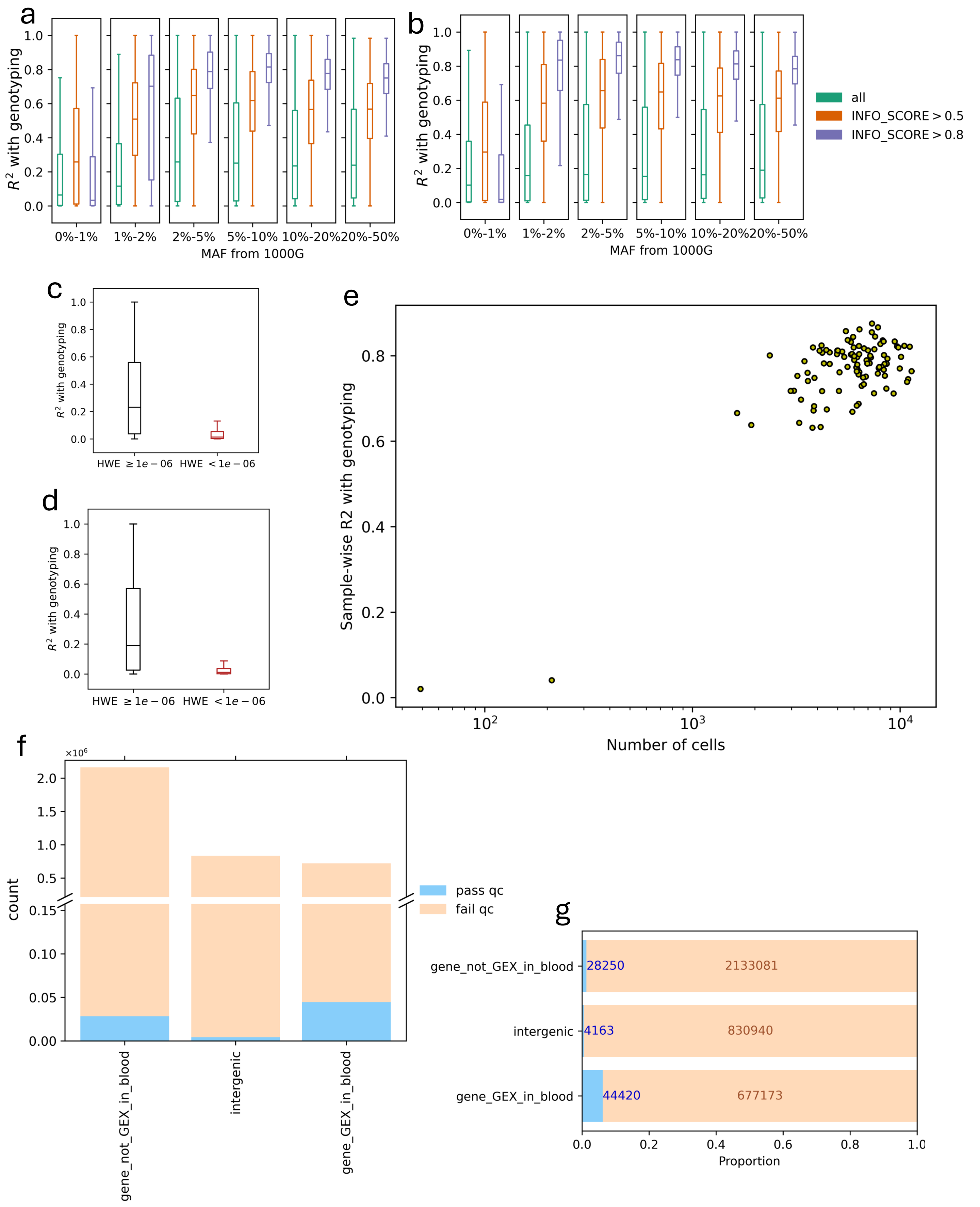


**Fig S1.** Genoype dosage concordance across minor allele-frequency (MAF) bins calculated using 1000 Genomes Project and INFO_SCORE filtering thresholds for GSA variants (**a**) and TOPMed-imputed variants (**b**). Genotype concordance above and below HWE threshold for GSA variants (**c**) and TOPMed-imputed variants (**d**). Sample-wise genotype concordance across number of cells (**e**). Number (**f**) and proportion (**g**) of variants annotated to genes that are expressed or not, or otherwise, that pass or fail quality control.

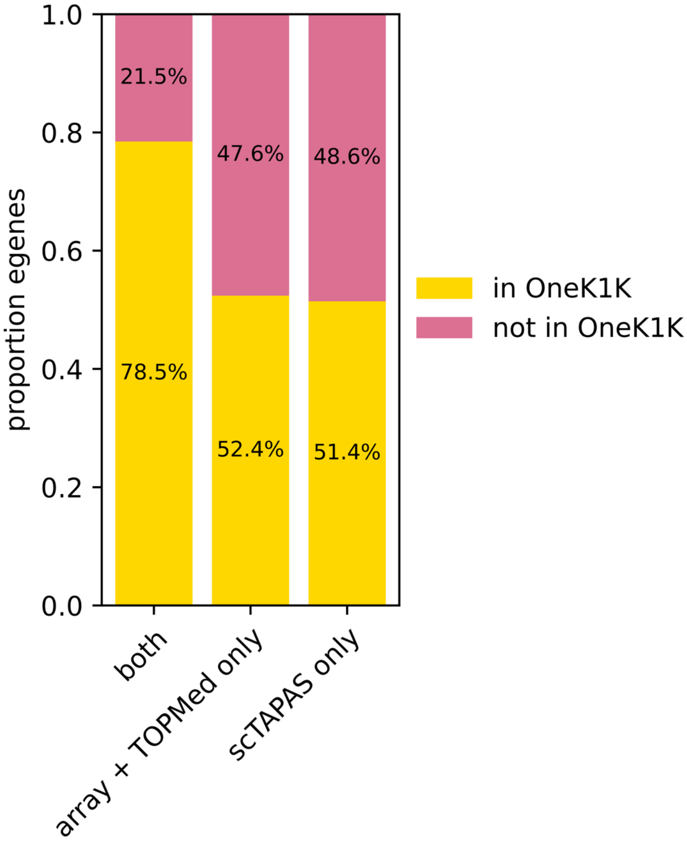


**Fig S2.** Proportion of eGenes overlapping with OneK1K for at least one matching cell type

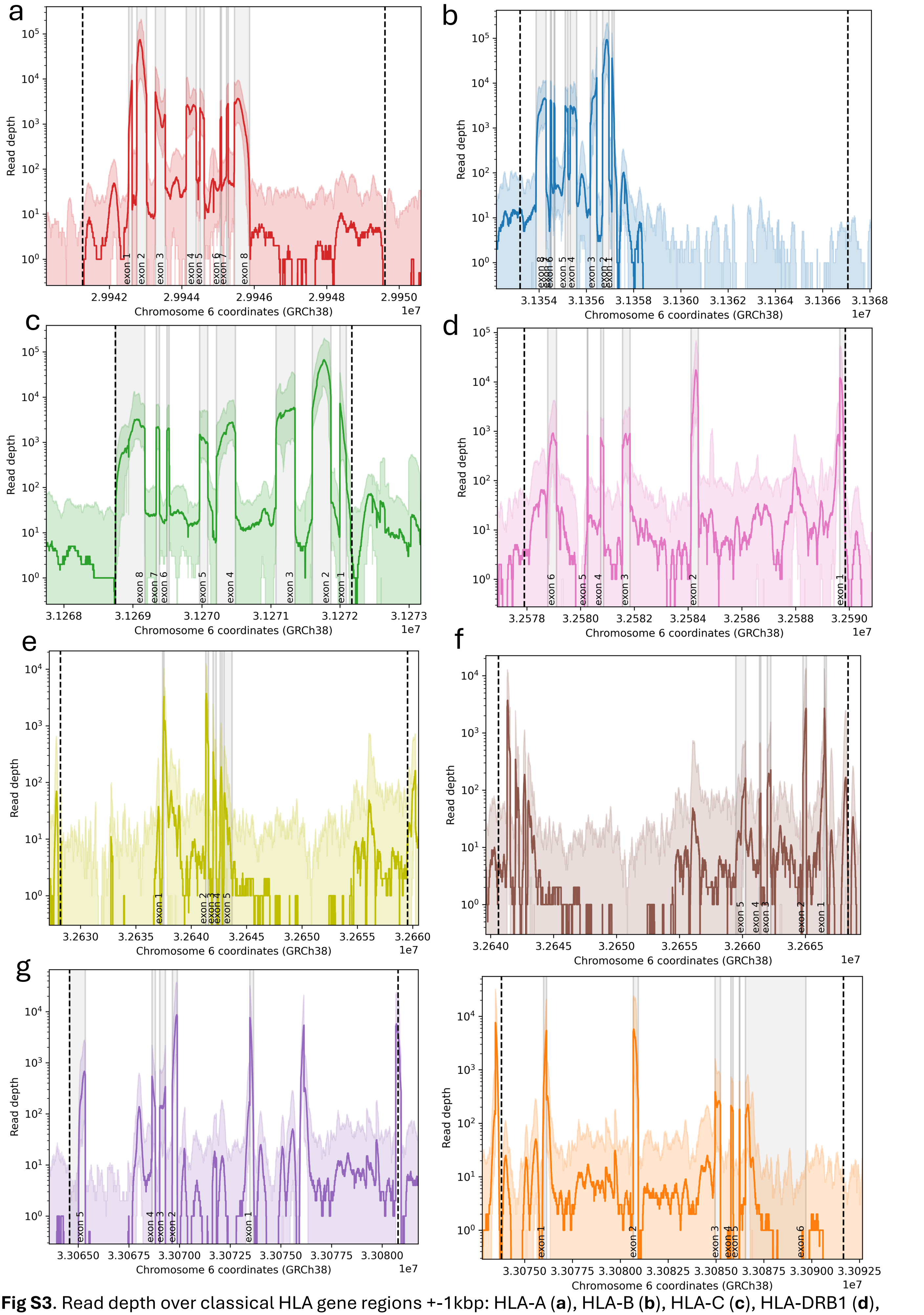


**Fig S3**. Read depth over classical HLA gene regions +-1kbp: HLA-A (**a**), HLA-B (**b**), HLA-C (**c**), HLA-DRB1 (**d**), HLA-DQA1 (**e**), HLA-DQB1 (**f**), HLA-DPA1 (**g**), HLA-DPB1 (**h**). Median shown in bold, minimum and maximum filled above and below.

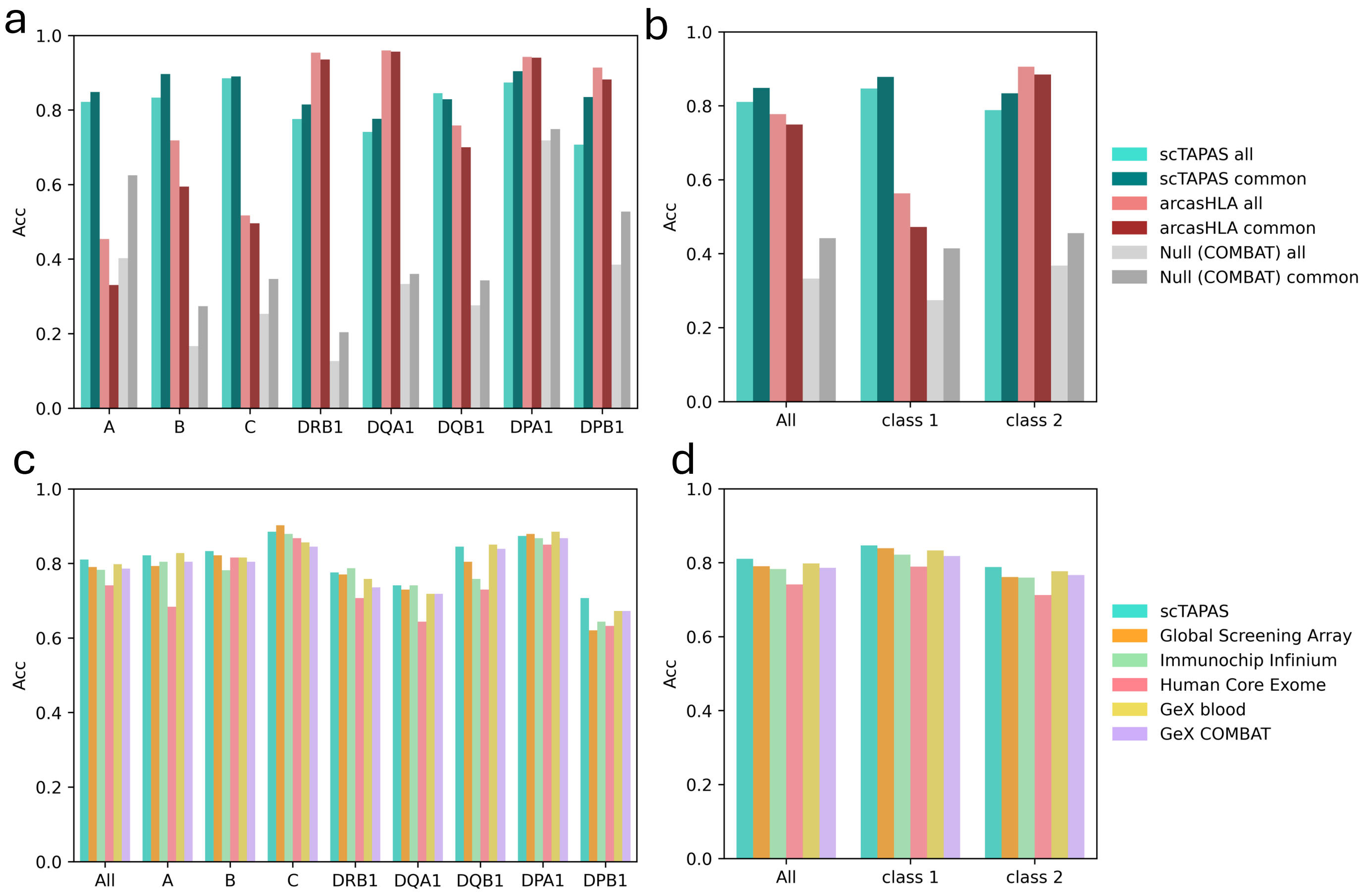


**Fig S4**. HLA gene-wise (**a**) and summarised gene-wise (**b**) accuracy for scTAPAS, arcasHLA and null model for all and common HLA alleles at one-field. HLA gene-wise (**c**) and summarised gene-wise (**d**) accuracy for scTAPAS and variations of scTAPAS where subsets of variants are extracted prior to HLA imputation.
